# Characterization of Genomic Diversity In Bacteriophages Infecting *Rhodococcus*

**DOI:** 10.1101/2022.11.23.517428

**Authors:** Dominic R. Garza, Daria Di Blasi, James A. Bruns, Brianna Empson, Isabel Light, Maisam Ghannam, Salvador Castillo, Britney Quijada, Michelle Zorawik, Ana E. Garcia-Vedrenne, Amanda C. Freise

## Abstract

Bacteriophages are globally ubiquitous viruses that infect bacteria. With nearly 4,000 sequenced genomes of phages infecting the phylum Actinobacteria available, genomic analyses of these actinobacteriophage genomes has been instrumental in uncovering a diverse genomic landscape often characterized by genome mosaicism. Here, we describe the genomic characterization of 57 sequenced phages capable of infecting the genus *Rhodococcus.* These phages were previously isolated at multiple institutions by students in the SEA-PHAGES program using four different species of *Rhodococcus.* Most *Rhodococcus* phages have been grouped into 4 clusters based on their genomic similarities; 13 phages are singletons too genetically distinct for clustering. These clusters and singletons contain *Siphoviridae* and *Myoviridae* phages, and most contain integrase and repressor genes indicative of a potential lysogenic life cycle. The genome size of these phages varies from 14,270 bp to 142,586 bp and their G+C% content ranges from 41.2–68.4%, while that of their *Rhodococcus* hosts typically exceeds 60%. Through comparative genomic analyses, it was revealed that these *Rhodococcus* phages display high intracluster similarity but low intercluster similarity, despite their shared ability to infect the same host genus. Additionally, these *Rhodococcus* phages share similarities with phages that infect other Actinobacterial hosts such as *Gordonia, Streptomyces* and *Arthrobacter.*

## Introduction

Bacteriophages, or phages, are globally ubiquitous viruses capable of infecting bacteria. They account for an overwhelming portion of the biosphere’s entities and their genomes represent a wealth of genetic diversity (Hatfull & Hendrix, 2011). Actinobacteriophages, phages capable of infecting Actinobacteria, have been of particular interest to researchers in the past 20 years due to their relevance as potential solutions to a variety of issues such as antibiotic resistance, food contamination and more (Sohail et al., 2020).

Currently, actinobacteriophages have been isolated on a variety of bacterial host strains belonging to 14 different genera, with a majority of these phages infecting the genera of *Mycobacterium, Gordonia, Microbacterium* or *Arthrobacter* (Hatfull, 2020). Of the over 20,000 actinobacteriophages that have been isolated to date, nearly 4,000 have been sequenced and their genomes made available in Genbank. While the number of isolated and sequenced actinobacteriophages has increased dramatically in recent years, much of their genomic landscape remains largely unexplored and more in-depth analyses of their genomes are needed to better understand their impact on the world around us (Hatfull, 2015). Thus, characterizing the genomic diversity of actinobacteriophages has been of great interest to the scientific community.

Investigations into the mycobacteriophage population, accounting for the majority of all isolated and sequenced actinobacteriophages, has revealed an immense realm of genetic diversity. These mycobacteriophage genomes are mosaic in structure with their genomes constructed of novel combinations of gene modules exchangeable with other phages (Hatfull, 2008). Additionally, their genomes constitute a continuum of diversity fueled by the delivery of genes from sources outside the mycobacteriophages themselves (Pope et al., 2015). Other investigations into phages infecting the genera *Gordonia* (Pope et al., 2017), *Arthrobacter* (Klyczek et al., 2017) and *Microbacterium* (Jacobs-Sera et al., 2020) have revealed similar trends in phage diversity. With large-scale comparative genomic analyses already performed on the most-sampled genera of actinobacteriophages, there is still the need to perform similar analyses on phages infecting other well-sampled genera such as *Streptomyces* or *Rhodococcus.*

To expand on the current knowledge of genetic diversity among actinobacteriophages, we investigated phages isolated from environmental samples using the host genus of *Rhodococcus.* The *Rhodococcus* genus comprises Gram-positive bacteria containing mycolic acids in their cell wall (Bell et al., 1998). These bacteria are commonly found in soil, water and in eukaryotic cells (McLeod et al., 2006). *Rhodococcus* has played a critical role as a biocatalyst in the synthesis of several organic compounds such as acrylamide, used in the production of polyacrylamide for applications such as oil recovery and water treatment (Jiao et al., 2020). Additionally, there has been some success in the application of *Rhodococcus* for bioremediation efforts in contaminated environments such as air, water and soil (Kuyukina & Ivshina, 2010). Currently, there is only a single known pathogenic species of *Rhodococcus, R. equi,* that has been the causative agent of pneumonia in young horses, immunocompromised horses and immunocompromised humans (Stewart et al., 2019). Therefore, the role of *Rhodococcus* species in bioproduction, bioremediation and disease highlights the need for a better general understanding of *Rhodococcus* phages.

As of November 2022, only 30 *Rhodococcus* phages had been described in the literature (Bonilla et al., 2017; Petrovski et al., 2012, 2013a, 2013b; Salifu et al., 2013; Summer et al., 2011). Here, we build upon these previous analyses to examine the general diversity of *Rhodococcus* phage genomes. We describe the isolation of 126 phages infecting the genus *Rhodococcus.* Most known *Rhodococcus* phages have been discovered through the efforts of students belonging to large-scale inclusive Research Education Communities (iRECs) such as the Science Education Alliance-Phage Hunters Advancing Genomics and Evolutionary Science (SEA-PHAGES) program and the Phage Hunting Integrating Research and Education (PHIRE) program (Hanauer et al., 2017; Russell & Hatfull, 2017). The *Rhodococcus* phages discovered by members of the SEA-PHAGES or PHIRE programs were isolated on one of the following 4 *Rhodococcus* species: *R. equi, R. erythropolis, R. globerulus* and *R. rhodochrous.* Of these 126 *Rhodococcus* phages, 57 have been sequenced and manually annotated as of November 2022. Thus, we used these 57 phages to provide a general characterization of the *Rhodococcus* phage population’s genomic diversity. In general, phages that share considerable gene content similarity can be placed into groups with other phages called clusters. As of November 2022, *Rhodococcus* phages make up 4 different clusters – CA, CB, CC and CE – as well as 13 singletons, a category consisting of only one single phage that lacks any closely related relatives (Hatfull, 2020). Bioinformatic tools such as nucleotide and amino acid sequence similarity dot plots, gene content similarity (GCS) maps, visual genome maps and SplitsTree analyses were used to assess similarities and differences between and within the 4 *Rhodococcus* phage clusters and representative singletons. Through comparative genomic analyses, it was revealed that these *Rhodococcus* phages display high intracluster similarity but low intercluster similarity, despite their shared ability to infect the same host genus.

## Materials and Methods

### Bacterial Strains

All phages were isolated on one of four different species of *Rhodococcus: R. equi, R. erythropolis, R. globerulus* and *R. rhodochrous* (Supplemental Table 1). The following 6 strains of *R. equi* were used: *R. equi* 05-305*, R. equi* 05-306, *R. equi* HDP1C, *R. equi* MillB, *R. equi* NCIMB 10027 and *R. equi* Requ28. The following 3 strains of *R. erythropolis* were used: *R. erythropolis* NRRL B-1574*, R. erythropolis* Rery29 and *R. erythropolis* RIA 643. The following 2 strains of *R. globerulus* were used: *R. globerulus* NRRL B-16938 and *R. globerulus* Rglo35. For *R. rhodochrous,* only the single strain of *R. rhodochrous* Rrho39 (DSMZ43241) was used in this study.

### *Rhodococcus* Phage Isolation, Purification, Amplification, and Virion Analysis

The *Rhodococcus* phages were isolated from soil samples collected by researchers at different institutions (Supplemental File 1) using either an enriched or direct isolation protocol as described previously (Petrovski et al., 2012, 2013a, 2013b; Salifu et al., 2013; Summer et al., 2011). PYCa broth (containing per 1 liter volume: 1.0 g Yeast Extract, 15.0 g Peptone, 2.5 ml of 40% Dextrose, 2.5 ml of 1 M CaCl_2_, and 1 ml of 1000X CHX) was used for phage isolation, purification and amplification. Phages were purified via multiple rounds of plaque picking and plaque assays to ensure a clonal phage population. Phage titers and plaque morphology were recorded during each round of purification to ensure consistency was maintained. A high-titer phage lysate was generated by flooding plaque assay plates showing webbed-lysis with phage buffer (containing per 1 liter volume: 10 ml of 1 M Tris stock (pH 7.5), 10 ml of 1 M MgSO_4_, 4 g NaCl, 10 ml of 100 mM CaCl_2_), incubating for at least 24 hrs and then filtering through a 0.22 μm filter. Using the collected high-titer phage lysates, each phage’s DNA was isolated, run on an agarose gel and assessed for purity as described previously (Summer, 2009).

### Genome Sequencing, Assembly and Annotation

For all *Rhodococcus* phages, 1 of 3 shotgun sequencing methods – Illumina, Ion Torrent, and Roche 454 – was used as described previously (Russell, 2018). Open Newbler (DS De Novo Assembler) on Linux software was used to establish a new Assembly Project and genome FASTA files were obtained.

Phage genomes were auto-annotated using DNA Master (Lawrence, 2007). GLIMMER (Delcher et al., 1999) and GeneMark (Besemer & Borodovsky, 2005) predicted protein-coding regions with respective start and stop sites. GeneMark coding potential maps were used to manually identify regions with reasonable coding potential (Besemer & Borodovsky, 2005). The Phamerator database grouped phage genes into phamilies (phams) of closely related proteins using pairwise sequence comparisons (Cresawn et al., 2011). Synteny was assessed using Phamerator maps to determine the likelihood of predicted genes being real. Suggested start sites were confirmed or rejected upon manual annotation. The Starterator database was used to compare auto-annotated start sites to the manually annotated start sites of other genes belonging to the same pham (Russell & Hatfull, 2017). Z-score and ribosome binding sites (RBS) were used to help to confirm these findings.

The function of each annotated gene was determined using PhagesDB BLASTp (https://phagesdb.org/blastp/), NCBI BLAST (Altschul et al., 1990), HHpred (Söding et al., 2005) and CDD (Marchler-Bauer et al., 2005). These tools compared the query protein sequence and structure to other known protein sequences and structures to predict the function of these genes. TOPCONS (Bernsel et al., 2009) and TMHMM (Krogh et al., 2001) were used to determine whether transmembrane domains (TMDs) were predicted in each gene’s protein sequence.

### Selection of Representative *Rhodococcus* Phages for Analysis

A gene content network phylogeny was created using PhamNexus and SplitsTree4 modeling in order to visualize intracluster and intercluster relationships based on shared gene content (Huson & Bryant, 2006). Using a Linux based virtual desktop (VirtualBox), pham data from all *Rhodococcus* phages were combined into a single nexus file using PhamNexus. The Nexus file was then uploaded onto the SplitsTree4 software application for iOS to render an unrooted tree using equal angle and Neighbornet algorithms.

Due to the disproportionately large size of Cluster CA compared to other *Rhodococcus* clusters, the generated SplitsTree model was used to select a smaller subset of phages that would best serve as representative of Cluster CA as a whole in later analyses. The representative phages were selected so that at least 1 phage from each grouping on the Splitstree model was included. The following 8 phages from Cluster CA were selected: AngryOrchard, CosmicSans, Espica, Krishelle, RER2, RGL3, Takoda and Yogi. All phages from Cluster CB (Grayson, Peregrin, Weasels2), CC (Pepy6, Poco6) and CE (NiceHouse, Trina) were selected for further analysis. Ultimately, 6 *Rhodococcus* singletons – E3, Finch, Jace, REQ2, REQ3 and Whack – were selected for analysis since they formed pairs with one another where their shared gene content similarity (GCS) exceeded at least 13.7%. The 8 other singletons showed very limited similarity to all other *Rhodococcus* phages.

### Nucleotide and Amino Acid Sequence Similarity

In order to provide a holistic characterization of the *Rhodococcus* phages, their nucleotide and amino acid sequence similarities were assessed using Genome Pair Rapid Dotter (GEPARD) (v1.40) (Krumsiek et al., 2007), a software capable of generating dotplots to highlight similarities and differences between sequences. As previously mentioned, the 8 representative Cluster CA phages and 6 singletons were selected for analysis. All phages from Clusters CB, CC and CE were included as well.

FASTA files for the whole-genome nucleotide sequences of these phages were obtained from PhagesDB (Russell & Hatfull, 2017) and merged into a single .txt file. Concatenated FASTA files of complete proteome amino acid sequences found on Genbank (Bilofsky et al., 1986) were utilized for amino acid comparisons. Word sizes of 10 and 5 were implemented as a similarity threshold for nucleotide and amino acid similarity, respectively. Rendered dot plots were combined using the Crop Overlay feature on Microsoft PowerPoint in order to depict nucleotide and amino acid data within one image.

### Gene Content Similarity (GCS)

GCS calculations were visualized in a heat map to assess the abundance of shared phams among the *Rhodococcus* phage clusters and selected singletons (Cresawn et al., 2011). GCS is a pairwise value and was calculated by finding the proportion of shared phams between two phages as a ratio of the number of total phams in each phage. The GCS is the average of these two values. The gene lists for all *Rhodococcus* phages used in this study were acquired from PhagesDB (Russell & Hatfull, 2017). GCS was computed using the GCS tool on PhagesDB (https://phagesdb.org/genecontent/) for all Cluster CA, CB, CC and CE phages and previously selected representative singletons E3, Finch, Jace, REQ2, REQ3 and Whack. GCS values were used to generate a heat map on Prism (www.graphpad.com).

### Phamerator

Synteny, shared phams, genome architecture, and nucleotide sequence similarity were compared using Phamerator (https://phamerator.org/) (Cresawn et al., 2011). 8 *Rhodococcus* phages (2 from each cluster) were selected for Phamerator analysis. The 2 phages selected from each of Cluster CA (RGL3 and Rhodalysa) and CB (Peregrin and Weasles2) were chosen for comparison as they displayed the lowest GCS values within their respective clusters, providing a conservative estimate of genomic similarities. All phages in Clusters CC and CE were chosen by default as both clusters contained only 2 phages.

### Evolutionary Relationships Between *Rhodococcus* Clusters and *Non-Rhodococcus* Phages

To investigate the relationship between *Rhodococcus* clusters and phages of other clusters, the whole-genome nucleotide sequence of all Cluster CB, CC and CE phages, as well as the 8 previously selected representative Cluster CA phages, were individually uploaded onto NCBI BLAST (Altschul et al., 1990) to identify similar nucleotide sequences. For each phage, the top 5 NCBI BLAST hits and 1-3 other hits from phages isolated on hosts other than *Rhodococcus* (e.g., *Gordonia, Mycobacterium, Streptomyces,* etc.) were selected for analysis. The GCS values between each *Rhodococcus* cluster – CA, CB, CC and CE – and those of the other phages selected from the NCBI BLAST results were calculated using the GCS tool on PhagesDB (https://phagesdb.org/genecontent/). Those *non-Rhodococcus* phages sharing over 15% GCS with at least one other *Rhodococcus* phage were selected for further analysis. The following *non-Rhodococcus* phages were included in this GCS analysis: MrMiyagi (AC), ElephantMan (AU1), Teacup (AU2), JimJam (BE2), TomSawyer (BE2), Manuel (BF), TaidaOne (BI1), Blueeyedbeauty (BK1), Stigma (BK1), SparkleGoddess (BK1), JustBecause (BM), Darwin (EN), MsGreen (L3), TyDawg (M1), Cosmo (V) and all 28 Cluster DJ phages.

Phamerator (https://phamerator.org/) (Cresawn et al., 2011) was used to align these phages’ genomes to assess synteny, shared pham content and overall genomic architecture. The Phamerator maps included only those non-*Rhodococcus* phages sharing the highest and lowest GCS values with the *Rhodococcus* clusters in order to better highlight the range of similarities and differences existing between these groups of phages. The following non-*Rhodococcus* phages were included in this Phamerator analysis: Pepy6 (CC), Poco6 (CC), Duffington (DJ) and Secretariat (DJ).

SplitsTree modeling was performed to offer a visual representation of those relationships identified between the *Rhodococcus* phages and other representative phages that infected either *Gordonia, Streptomyces* or *Arthrobacter.* The following *non-Rhodococcus* phages were included in this SplitsTree analysis: DBQu4n (A2), PurpleHaze (A3), Burger (A4), Dublin (A5), Philis (A8), RhynO (A10), Zimmer (A12), JSwag (A15), MyraDee (A18), ElephantMan (AU1), Teacup (AU2), JimJam (BE2), TaidaOne (BI1), Stigma (BK1), SparkleGoddess (BK1), Duffington (DJ) and Secretariat (DJ).

## Results and Discussion

### *Rhodococcus* phage isolation

As of November 2022, PhagesDB.org listed 126 phages isolated from environmental samples (soil, compost, manure and wastewater) using 4 different species of *Rhodococcus.* Most of these *Rhodococcus* phages were isolated using the species of *R. erythropolis* (116) with a smaller subset on *R. equi* (8), *R. globerulus* (1) and *R. rhodochrous* (1) (Supplemental Table 1). These *Rhodococcus* phages were isolated using either enrichment or direct plating.

Most of these 126 phages were isolated through the efforts of students participating in either the SEA-PHAGES or PHIRE programs at University of Maine – Honors College, University of Wisconsin - River Falls, University of Pittsburgh, Carnegie Mellon University, University of Louisiana at Monroe, Nyack College, College of William & Mary, Florida Gulf Coast University, Morehouse College, Cabrini University, Virginia Commonwealth University, Wilkes University, Texas A&M University, University of Edinburgh, and La Trobe University. Most of the phages were isolated from samples from across the United States of America, with only a few phages isolated from the United Kingdom or Australia. While these 126 phages account for a considerable portion of the isolated *Rhodococcus* phage population, other *Rhodococcus* phages have been discovered by members of the scientific community outside of the PHIRE and SEA-PHAGES programs on other species such as *R. australis* and *R. opacus* (Gill et al., 2018; Hiddema et al., 1985; *Rhodococcus Phage Mbo2, Complete Genome*, 2022; *Rhodococcus Phage Mbo4, Complete Genome,* 2022). However, these phages are not listed on PhagesDB.org and were subsequently not included in the analyses presented in this paper.

As of November 2022, 58 of the 126 isolated *Rhodococcus* phages have been sequenced. *Rhodococcus* phage genomes were sequenced at the Pittsburgh Bacteriophage Institute and the UCLA Genotyping and Sequencing Core. Following sequencing, each *Rhodococcus* phage’s genome was auto-annotated. Only 57 of the 58 sequenced phages have been manually annotated. Singleton TroggleHumper is currently being annotated by students at Queensborough Community College. Among those *Rhodococcus* phages that have been sequenced, about 50% of them have been found to produce clear plaques on their respective host species; however, 8 phages were recorded as capable of producing turbid plaques as well. Through the use of transmission electron microscopy (TEM), it was found that 49 of the 58 sequenced *Rhodococcus* phages are of the *Siphoviridae* morphotype. The only exceptions to this trend appear to be singleton phages E3 and Finch, both of which showcase the *Myoviridae* morphotype. As of November 2022, 7 sequenced *Rhodococcus* phages currently have no available TEM image to determine their respective morphotypes: PhailMary (CA), Shuman (CA), Swann (CA), Trina (CE), ChewyVIII (Singleton), Jace (Singleton) and Sleepyhead (Singleton).

### Genomic trends within and between the *Rhodococcus* clusters

*Rhodococcus* phages account for the sixth most sequenced group of phages infecting a single genus within the larger actinobacteriophage community. However, only a subset of Cluster CA phages have been previously described using comparative genomic analysis (Bonilla et al., 2017). In order to address this gap in knowledge, our research extends this same type of analysis to include all currently annotated phages. Each bioinformatic tool described below contributed to this effort by providing a detailed look into the intracluster relatedness, intercluster relatedness and evolutionary relationships of the 4 existing *Rhodococcus* clusters – CA, CB, CC and CE – and 6 representative singletons: E3, Finch, Jace, REQ2, REQ3 and Whack. These singletons were selected for analysis since they formed pairs with one another where their shared gene content similarity (GCS) exceeded at least 13.7%. The 8 other singletons were omitted since they showed very limited similarity to all other *Rhodococcus* phages.

### General trends of intracluster and intercluster relatedness across the *Rhodococcus* clusters

Investigations into the relatedness within and among the *Rhodococcus* clusters began with a SplitsTree model. The SplitsTree was used to visualize a gene content-based network to assess relationships of shared phams across the sequenced *Rhodococcus* phages (Figure 1). The SplitsTree analysis revealed that phages in each *Rhodococcus* cluster were distinct from phages of other *Rhodococcus* clusters. Within Cluster CA, most phages had very short branches between one another, such as Naiad and Natosaleda, which indicated very high intracluster relatedness (Figure 1). However, phage RGL3 represented an outlier to this trend, as it displayed a relatively longer branch which indicated low levels of similarity to the other CA phages. Cluster CC phages were connected by relatively short branch lengths as well, indicating high levels of intracluster relatedness (Figure 1). It should be noted that Cluster CB and CE phages displayed low intracluster similarities relative to clusters CA and CC as indicated by their long connecting branches (Figure 1). Cluster CA and CC phages shared a divergence point from phages belonging to Clusters CB and CE. Cluster CC phages Pepy6 and Poco6 displayed the shortest branch distance from Cluster CA phages, suggesting a higher degree of similarity between the two clusters.

**Fig 1.**
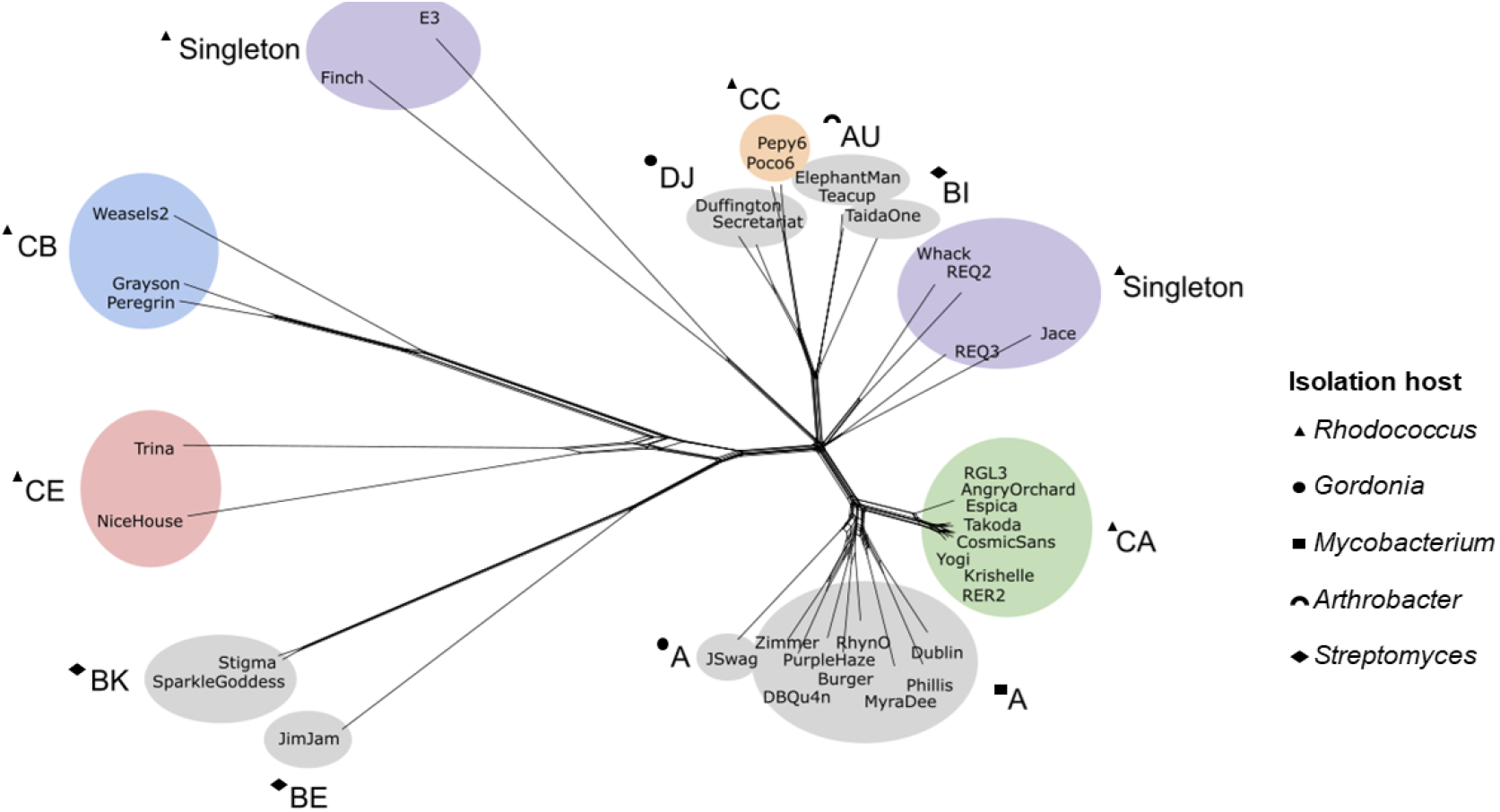
SplitsTree representation of relationships among *Rhodococcus* phages and select *non-Rhodococcus* phages. SplitsTree4 was used to visualize phage relationships based on shared phams. Clusters based on GCS are color coded; isolation host is denoted by a symbol. Gray overlays indicate *non-Rhodococcus* clusters. The scale bar indicates .01 substitutions per site. *Rhodococcus* phages are diverse, as indicated by the variable branch distances between clusters and singletons. Some *Rhodococcus* phages are more similar to phages from *non-Rhodococcus* hosts. For example, Cluster CC phages appear to show a closer relationship to Cluster DJ *(Gordonia),* Cluster AU *(Arthrobacteŕ)* and Cluster BI *(Streptomyces)* phages than with other *Rhodococcus* phages.

### Whole-genome sequence comparisons within and between *Rhodococcus* clusters

The general trends of intracluster and intercluster relatedness between the *Rhodococcus* clusters were investigated further using sequence analyses of whole phage genomes at nucleotide and amino acid levels. This was achieved using the Genome Pair Rapid Dotter (GEPARD) software, a visualization tool that generates dotplots to highlight similarities and differences between the provided nucleotide or amino acid sequences (Krumsiek et al., 2007). Nucleotide sequence similarity was investigated to provide a comparative understanding of the *Rhodococcus* phage genomes at the most basic level, while amino acid sequence similarity was employed to provide context for those evolutionary relationships reflected at the protein level (Figure 2).

**Fig 2.**
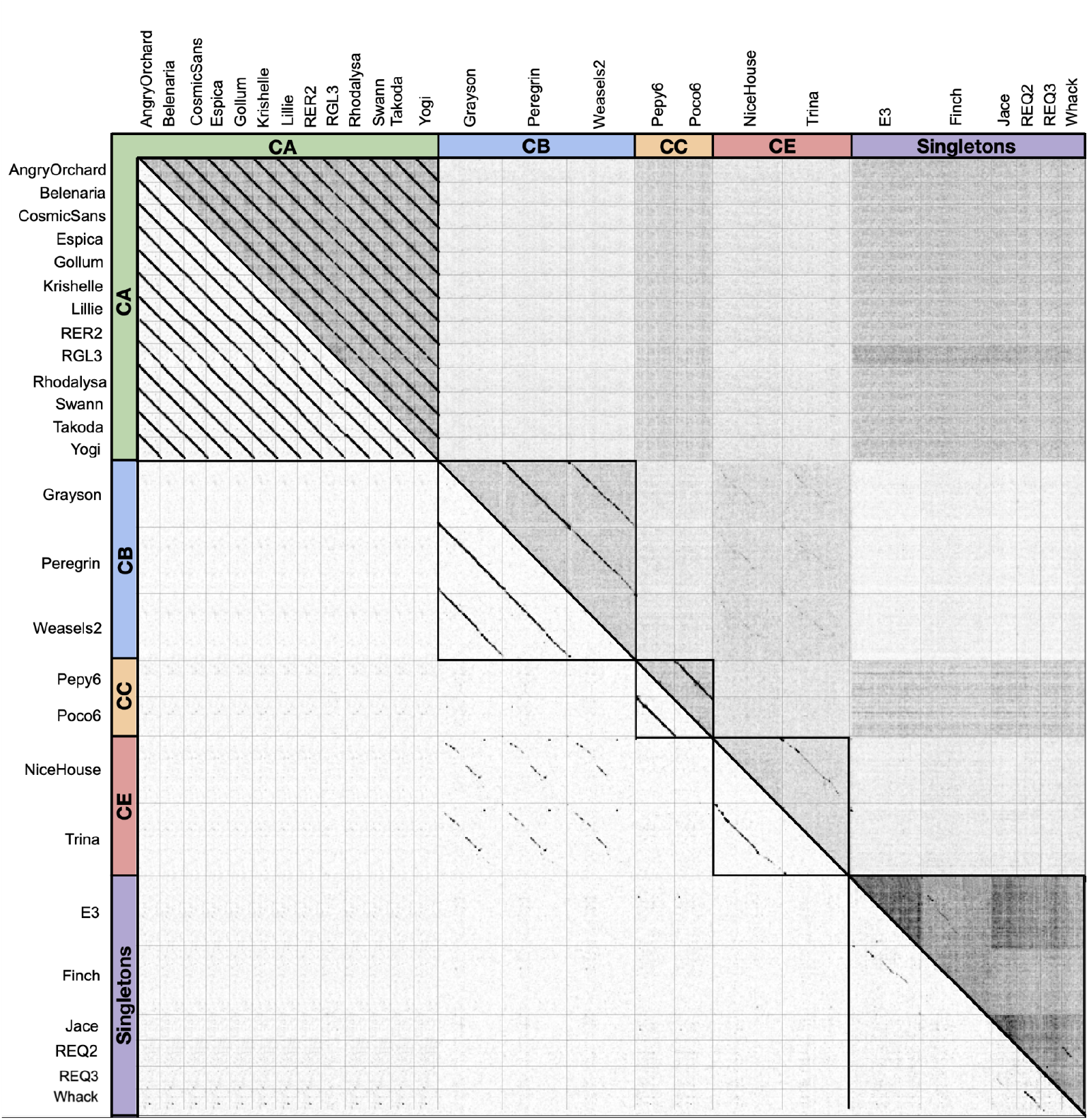
Dot plot of nucleotide and amino acid sequence similarity between 26 representative *Rhodococcus* phages’ complete genome sequences. The whole-genome nucleotide and complete proteome amino acid sequences for the 26 representative *Rhodococcus* phages were each compiled into a single text file in FASTA format and uploaded to Genome Pair Rapid Dotter (GEPARD) (Krumsiek et al., 2007). Dot plots were overlaid on top of each other, with nucleotide data on the top right half and amino acid on the bottom left half. For nucleotide sequence similarity comparisons, the word size was set to 10. For amino acid sequence similarity comparisons, the word size was set to 5. All *Rhodococcus* phage clusters display stronger intracluster similarity in regard to both nucleotide and amino acid sequence similarity when compared against intercluster similarity. Cluster CA phages display the strongest intracluster similarity for both nucleotide and amino acid sequence while Cluster CB and CE phages share the strongest intercluster similarity for both sequence types.

Cluster CA phages displayed the highest levels of intracluster relatedness relative to the other *Rhodococcus* clusters for both nucleotide and amino acid sequence comparisons. However, these Cluster CA phages showed low levels of intercluster similarity for both levels of sequencing when compared against the other *Rhodococcus* clusters (Figure 2). Similarly, Cluster CC phages also displayed high levels of intracluster relatedness and low levels of intercluster relatedness with respect to both nucleotide and amino acid sequence comparisons. The high degree of similarity between Cluster CC phages Pepy6 and Poco6 with Cluster CA phages highlighted by the SplitsTree model was less pronounced at both sequence levels, as the Cluster CA phages were shown to be well-defined and distinct from the other *Rhodococcus* clusters (Figure 1, Figure 2). Both Cluster CB and CE phages showed a similar trend to Cluster CC phages; however, both clusters’ levels of intracluster similarity were relatively lower than that of Clusters CA and CC. Cluster CB and CE phages displayed the highest levels of intercluster similarity for both nucleotide and amino acid sequence similarities despite having the longest average genomes among the *Rhodococcus* clusters.

### Gene content similarity within and between *Rhodococcus* clusters

To investigate whether gene content trends within and between *Rhodococcus* clusters reflected the same patterns as was revealed by whole-genome sequence analyses, the metric of gene content similarity (GCS) was employed. GCS was calculated between all phages belonging to the four *Rhodococcus* clusters as well as the 6 previously selected representative singletons. (Figure 3).

**Fig 3.**
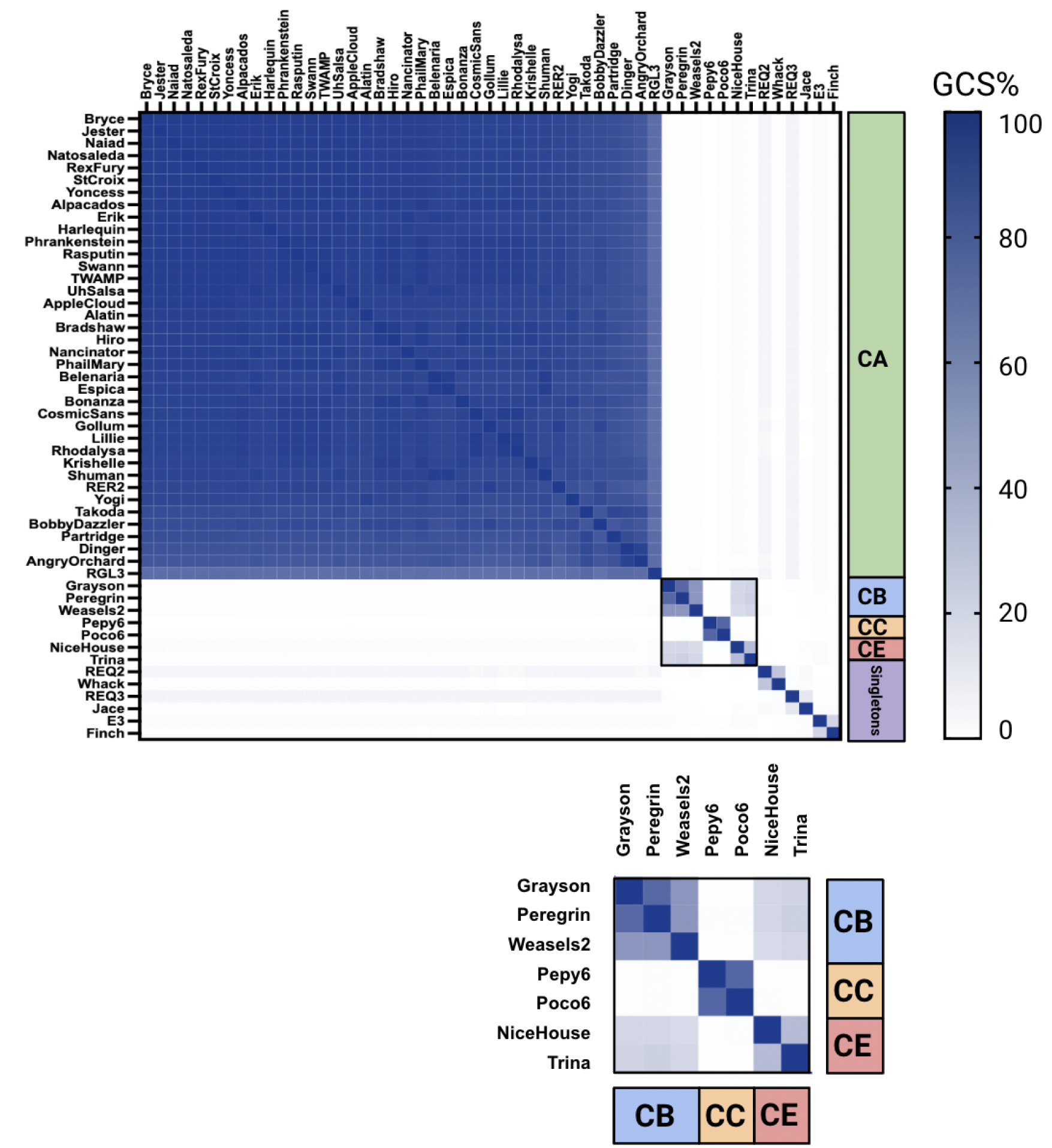
Heat map of the gene content similarity (GCS) among *Rhodococcus* phages. The PhagesDB Gene Content tool was used to calculate GCS values between 51 sequenced *Rhodococcus* phage genomes. Singletons sharing GCS values under 18.9% with all other *Rhodococcus* phages were excluded from this figure. Prism was used to generate a heat map representation of these GCS values. Each GCS value is represented by its coloring where lighter colored entries indicate lower GCS and darker shades depict higher GCS. There are high levels of intracluster GCS and low levels of intercluster GCS across *Rhodococcus* phage clusters CA, CB, CC and CE.

Cluster CA phages displayed the highest levels of intracluster similarity with all their GCS values exceeding 75%. There were even 17 Cluster CA phages that shared GCS values of 100% with one other CA phage. Cluster CB phages display GCS values greater than 58% for all phages and Cluster CC phages display GCS values of 81.3%. Cluster CE phages displayed the lowest levels of intracluster similarity with GCS values of only 38.3% – just barely above the minimum 35% shared GCS required for clustering (Figure 3).

Contrary to the trend observed with intracluster similarity, the intercluster similarity of the *Rhodococcus* phages appeared to be low despite their ability to infect the same host genus. The intercluster GCS values among *Rhodococcus* phages consistently fell below 2%; however, Clusters CB and CE deviated from this trend as they had intercluster GCS values of up to 20% (Figure 3). Both of these clusters share the longest average genomes out of the *Rhodococcus* clusters (Figure 4). The isolated nature of the *Rhodococcus* clusters is reminiscent of a previous analysis performed on 46 *Arthrobacter* phages where it was found that 6 of the 9 *Arthrobacter* clusters had less than 10% of their genes shared by at least one other *Arthrobacter* cluster (Klyczek et al., 2017). Similar to phages that infect either *Mycobacterium, Gordonia* or *Microbacterium,* it is likely that the *Rhodococcus* clusters could become less discrete as more *Rhodococcus* phages are sequenced and reach numbers comparable to these other host types (Jacobs-Sera et al., 2020; Pope et al., 2015, 2017).

**Fig 4.**
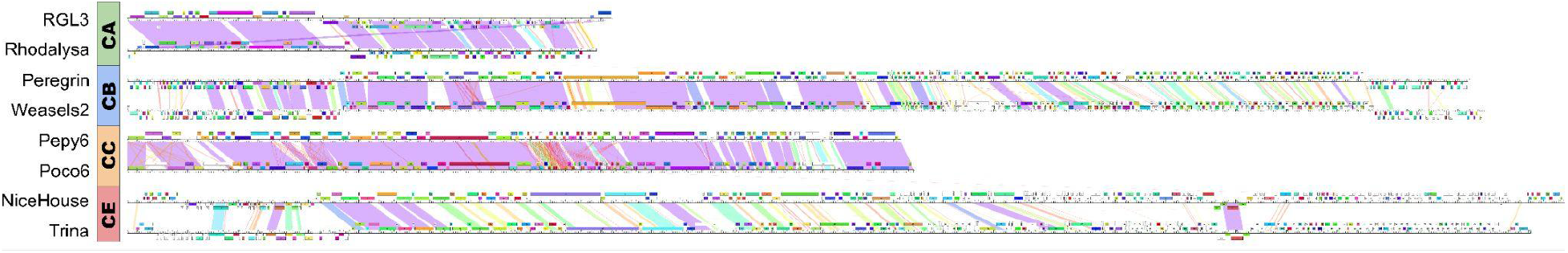
Genome comparison of *Rhodococcus* phage clusters CA, CB, CC and CE. Genome maps were made using Phamerator. Full genome comparison of phages RLG3 (CA), Rhodylsa (CA), Peregrin (CB), Weasles2 (CB), Pepy6 (CC), Poco6 (CC), NiceHouse (CE), and Trina (CE) shows high intra-cluster similarity and weak inter-cluster similarity. Each white ruler corresponds to an individual phage genome with the genes represented by colored boxes. Each gene’s color indicates its gene phamily (pham); white boxes represent orphams. The placement of the genes above and below the white rulers indicates rightwards and leftwards transcription, respectively. The shading between phages represents the degree of pairwise nucleotide sequence similarity between two phage genomes with purple indicating the highest similarity and red indicating lowest similarity.

### Distribution of phams across the *Rhodococcus* clusters

In order to localize the shared phams identified through GCS analysis, Phamerator was employed to examine synteny among *Rhodococcus* phages. Phamerator is a computational tool that allows for the visualization of genome architectures to assess nucleotide sequence similarity, shared phams and synteny among those selected phages (Cresawn et al., 2011).

The Phamerator maps confirmed prior investigations into the nucleotide sequence similarities of *Rhodococcus* phages using GEPARD dot plots, as phages from all 4 *Rhodococcus* clusters showed high levels of intracluster nucleotide sequence similarity (Figure 4). Additionally, for each of the 2 phages compared within a cluster, there were strong levels of synteny observed between phage genomes. Cluster CE phages showed many shared phams with one another despite showing relatively low levels of nucleotide sequence similarity (Figure 4). When comparing *Rhodococcus* phages of different clusters, there were little to no phams shared and nucleotide sequence similarity levels were sparse. Despite this, phages from Clusters CB and CE shared multiple phams and moderate levels of nucleotide sequence similarity (Figure 4).

In addition, a Phamerator map of those 17 CA phages sharing 100% GCS with one other Cluster CA phage (not shown) confirmed that the genome architecture of these phages was well conserved. These 17 CA phages included Rasputin, Alpacados, Espica, Belenaria, Lillie, CosmicSans, Rhodalysa, UhSalsa, Erik, Yoncess, Jester, Natosaleda, Naiad, RexFury, StCroix, Swann and Phrankenstein. The left arm of their genomes encodes structural genes (e.g., major capsid protein, major tail protein, and head-to-tail adaptor) while the right arm of their genomes encodes non-structural genes (e.g., DNA primase, DNA polymerase I, and ribonucleotide reductase). For these CA phages, both nucleotide sequence similarity and synteny were strong with the exception of a single gene: Alpacados_39. Across these 17 phages, this gene either belonged to one of two different phams or was completely absent from this area of the genome. As of now, there is no known function associated with this gene for either of the two phams. Phages Natosaleda, Naiad, RexFury and StCroix were the only genomes lacking this gene of the 17 CA phages.

### *Rhodococcus* phage clusters and singletons

The earliest *Rhodococcus* phages to be sequenced were completed back in the year 2011. Even though some *Rhodococcus* phages have been sequenced as recently as the year 2021, cluster membership for these 58 sequenced *Rhodococcus* phage genomes is still in compliance with the most recent clustering criteria of 35% shared gene content similarity (GCS). As of November 2022, 45 of the 58 sequenced *Rhodococcus* phages belong to one of the 4 *Rhodococcus* clusters. 13 phages do not meet the threshold of 35% shared GCS required for cluster membership, so these phages are currently designated as singletons. This proportion of clusters and singletons deviates from previous analyses that employed a similar quantity of phages exclusively infecting the host types of either *Mycobacterium* (60 phages), *Arthrobacter* (47 phages) or *Gordonia* (79 phages) (Hatfull et al., 2010; Klyczek et al., 2017; Pope et al., 2017).

#### Cluster CA

The majority of annotated *Rhodococcus* phages belong to Cluster CA (38 out of 57). There are currently no subclusters associated with Cluster CA. Most of these phages were isolated on the same host species of *R. erythropolis* (Supplemental Table 1). The only exception to this trend is phage RGL3, which was isolated on *R. globerulus* Rglo35. Of the 37 CA phages isolated on *R. erythropolis,* 34 were specifically isolated on the strain *R. erythropolis* RIA 643. The other 3 phages were isolated on either *R. erythropolis* NRRL B-1574, *R. erythropolis* Rery29 or *R. erythropolis* NRRL B-1574.

Approximately 90% of the CA phages are of the *Siphoviridae* morphotype with icosahedral heads and non-contractile tails (Supplemental Table 1). The remaining 10% of phages’ morphologies have not yet been confirmed due to the lack of available transmission electron microscopy (TEM) images. Of the CA phages, about 90% of these phages code for the functions of both an integrase and repressor, suggesting these phages are likely temperate. As of November 2022, four Cluster CA phages (BobbyDazzler, Nancinator, RER2 and RGL3) have been annotated to code for the function of integrase but not a repressor. However, all 4 phages contain at least one gene that belongs to a pham with at least one other gene member that codes for the function of immunity repressor: BobbyDazzler_58, Nancinator_59, RER2_51 and RGL3_52. This is likely a result of the fact that these 4 CA phages’ genomes were sequenced and annotated prior to the discovery of the other pham members that code for the function of immunity repressor. Therefore, it is likely that these 4 CA phages are temperate as well.

In general, the 38 CA phages have similar genome lengths (mean 46,597 bp; range 45,614 bp to 48,072 bp) (Figure 5). These phages’ genomes are relatively short in comparison to the other *Rhodococcus* clusters, as the other clusters all have phages with genomes above at least 76,000 bp. The G+C% content is consistent across the CA phages (mean 58.8%; range 58-59%), with RGL3 being the exception to this trend with a G+C% content of 62.7% (Supplemental Table 1, Figure 5).

**Fig 5.**
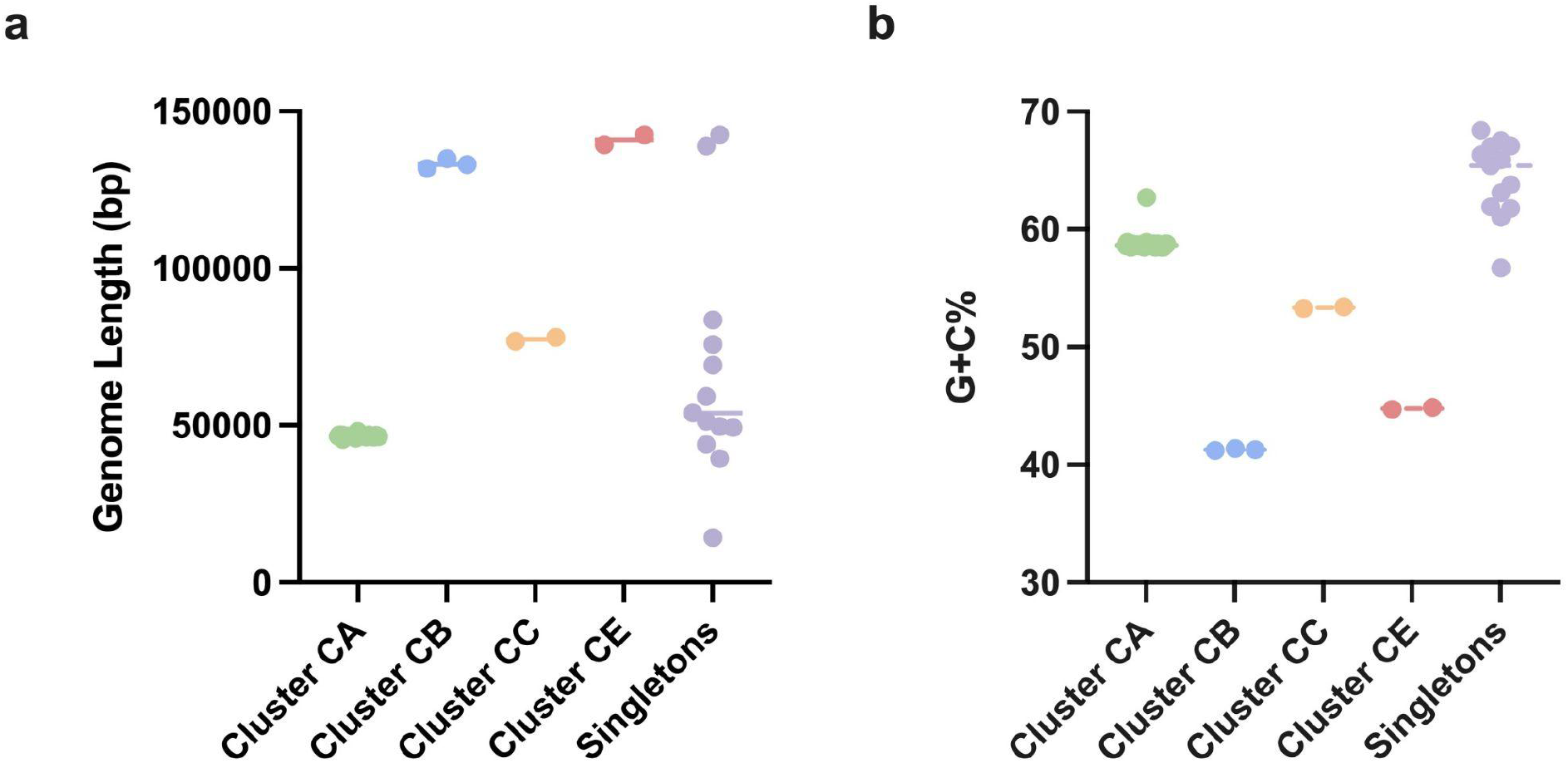
Average genome length (bp) and G+C% content of *Rhodococcus* phages. Prism was used to plot the **A.** genome length (bp) and **B.** G+C% content of all sequenced *Rhodococcus* phages. Dots represent individual phage genomes and horizontal lines represent the mean for that group. Cluster CA: n=38; Cluster CB: n=3; Cluster CC: n=2; Cluster CE: n=2; Singletons: n=13. Each line represents the average genome length (bp) of the phage cluster while each data point corresponds to an individual phage genome.

Comparisons of the aligned genomes demonstrate that the CA phages show extensive synteny (functionally similar genes occurring in the same relative order) (Figure 4). Their genomes are extremely similar to one another considering that all of their pair-wise shared gene content values exceed 75%. As previously mentioned, there are 17 Cluster CA phages that share 100% GCS with one other Cluster CA phage (Figure 3). The genome architecture of these CA phages is mostly conserved across Cluster CA members, with structural genes (e.g., major capsid protein, major tail protein, and head-to-tail adaptor) present in their genomes’ left arm and assembly genes (e.g., DNA primase, DNA polymerase I, and ribonucleotide reductase) in their genomes’ right arm.

For Cluster CA phages, the first half of their genomes is primarily rightwards transcribed with the second half of their genomes mostly leftwards transcribed (Figure 4). Some brief switches in transcription orientation do occur throughout their genomes. These switches in orientation are usually followed by only about 1-2 genes before switching back to the previous transcription orientation.

Orphams, genes lacking known close relatives, are relatively limited across the CA phage genomes. Out of 38 Cluster CA phages, only 10 phages contain at least 1 orpham in their genome. Most of these phages have 1-4 orphams present in their genomes, except RGL3 which has 11 orphams (Figure 4). RGL3 is the only sequenced phage isolated on *R. globerulus,* so this could account for the relatively higher percentage of orphams as compared with other CA phages that infect *R. erythropolis* instead.

#### Cluster CB

There are currently 3 Cluster CB phages (Grayson, Peregrin and Weasles2) that were isolated on the same host strain of *R. erythropolis* RIA 643 (Supplemental Table 1). The CB phages are all of the *Siphoviridae* morphotype with icosahedral heads. These phages have an average genome length of 133,260 bp (range: 131,801 bp to 134,973 bp) and average G+C% content of 41.3% (range: 41.2-41.4%) (Figure 5).

These CB phages have varying degrees of similarity to one another. Their average shared gene content is 65.93%; however, their pairwise shared gene content values range from 58.3% to 80.3% (Figure 3). Phages Grayson and Peregrin share 80.3% of their genes; the dissimilarity between their genomes becomes more pronounced in the downstream regions of their genomes’ right arm, where nearly half of the genes are orphams. Despite this, Grayson and Peregrin have a limited percentage of their genes that belong to orphams (6% and 7%, respectively). Weasels2 represents an outlier to this trend as nearly 30% of its genes belong to orphams. Weasles2 displays considerably less shared gene content with both Grayson and Peregrin; conserved regions between these phages are localized toward the middle of their genomes (Figure 4).

The Cluster CB phages have multiple switches between rightwards transcription and leftwards transcription across their genomes (Figure 4). These phages follow the same general pattern of transcription from the left to right arms of their genome: leftwards, rightwards, leftwards. Weasles2 deviates slightly from this trend, as its genome begins with a single rightwards transcribed orpham before following the same general pattern observed for the other Cluster CB phages. These 3 phages lack genes that have been associated with the functions of either integrase or repressor, and are likely lytic.

#### Cluster CC

There are currently 2 phages in Cluster CC: Pepy6 and Poco6. They are both capable of infecting the same host species of *R. equi* but Pepy6 and Poco6 were isolated on different strains (*R. equi* 05-306 and *R. equi* MillB, respectively). (Supplemental Table 1). Both phages are of the *Siphoviridae* morphotype with icosahedral heads. Their genome sizes are relatively similar to one another, but the genome of Poco6 (78,064 bp) is slightly longer than that of Pepy6 (76,797 bp). Poco6 and Pepy have genomes with similar G+C% content (53.3% and 53.4%, respectively) (Figure 5).

These phages have 77.57% shared gene content (Figure 3). Their shared gene content is not necessarily localized to a specific region of their genomes, as shared phams can be found across the entire length of their genomes (Figure 4). Those genes that are shared between the phages code primarily for functions related to structure/assembly (tail fiber protein, tape measure protein and head protein), genome packaging (portal protein, large terminase subunit protein and prohead protease) and lysis (holin, lysin A and lysin B). The synteny between this pair of phage genomes is discontinuous along their fully-aligned genomes (Figure 4). This disruption in synteny is accounted for mostly by orphams. The majority of these orphams appear at either end of their genomes with a smaller portion located in the middle (Figure 4). Pepy6 and Poco6 have 16 orphams and 13 orphams, respectively.

Neither of their genomes code for the functions of integrase or repressor which implies that Cluster CC phages are likely lytic. Their genomes are both exclusively rightwards transcribed (Figure 4).

#### Cluster CE

There are currently 2 phages that belong to Cluster CE: NiceHouse and Trina. They both infect the same host species of *R. erythropolis* but Nicehouse and Trina were isolated on different strains (*R. erythropolis* NRRL B-1574 and *R. erythropolis* RIA 643, respectively) (Supplemental Table 1). NiceHouse is of the *Siphoviridae* morphotype with an icosahedral head. The morphotype of phage Trina is currently undetermined because the transmission electron microscopy (TEM) image available is difficult to interpret due to image clarity (Holder et al., 2015).

NiceHouse and Trina have similar genome lengths (142,586 bp and 139,262 bp, respectively) that are considerably larger than the average genome lengths of Clusters CA and CC but similar to that of Cluster CB (Figure 5). These phages display only 37.5% shared gene content, just barely above the 35% threshold required for clustering (Pope et al., 2017) (Figure 3). About 40% of the genes present in each genome are orphams. The majority of their shared genes are localized to the center of their genomes (Figure 4).

Although these phages share only 37.5% of their genes, nucleotide sequence similarity between their genomes is sparse across the entire length of their genomes (Figure 2). Patches of nucleotide sequence similarity appear throughout their genomes but span only 1-3 genes (Figure 4). Their shared genes primarily code for functions related to virion structure (e.g., major capsid protein or minor tail protein), assembly (e.g., tail assembly chaperone or portal protein) and lysis (e.g., lysin B or hydrolase).

These phages’ genomes have multiple switches in the direction of their transcription (Figure 4). Both phages follow a similar transcription pattern from the left to right arms of their genome – rightwards, leftwards, rightwards, leftwards and rightwards – when it comes to switches in the orientation of their genomes’ transcription. Their genomes lack any genes associated with the functions of either an integrase or repressor, so it is likely that these CE phages are lytic.

#### *Rhodococcus* Singletons

Of the 58 sequenced *Rhodococcus* phages available on PhagesDB.org, 13 have been designated as singletons. These singleton phages are too genetically distinct to warrant either inclusion into current *Rhodococcus* clusters or the formation of a new cluster. The *Rhodococcus* singletons account for roughly 22% of the total Actinobacteriophage singleton population, followed by *Streptomyces* singletons (19%), *Microbacterium* singletons (14%), *Mycobacterium* singletons (11%) and *Gordonia* singletons (11%). In general, the *Rhodococcus* singletons share pairwise GCS values between 0-18.9% with all other *Rhodococcus* phages. However, two pairings of singletons display pairwise GCS values outside this range: REQ2 with Whack (30.2%) and E3 with Finch (22.33%) (Figure 3). Here we combine these similar singletons for analysis, leaving the other 8 singletons to be investigated individually. Singleton TroggleHumper was omitted from our analyses since its genome is currently being annotated by students at Queensborough Community College.

#### Singletons REQ2 and Whack

These singletons share a pairwise GCS value of 30.2% with one another and display GCS values of less than 13% with all other *Rhodococcus* phages (Figure 3). REQ2 and Whack were isolated on different host species *(Rhodococcus equi* Requ28 and *Rhodococcus erythropolis* NRRL B-1574, respectively) (Supplemental Table 1). Both phages are of the *Siphoviridae* morphotype. REQ2 and Whack have similar genome lengths (49,330 bp and 49,660 bp, respectively) but differ slightly in terms of their genomic G+C% content (65.4% and 61.9%, respectively).

A few common features are shared between the overall genomic structure of these phages (not shown). REQ2 and Whack show similar transcription orientation patterns across their genomes. The ends of both phages’ genomes are rightwards transcribed with a small block of genes that are leftwards transcribed located in the middle of their genomes. The majority of those genes shared between these phages can be localized to the left arm of their genome that is rightwards transcribed. Most of the genes shared between REQ2 and Whack code for functions related to virion structure (e.g., major/minor tail proteins, major capsid protein, and portal protein). For both phages, approximately 40% of their genes belong to orphams. In general, most of these orphams are localized to the right arm of their genomes. Both REQ2 and Whack code for the functions of integrase and repressor, suggesting that these phages may both be temperate (Supplemental Table 1).

#### Singletons E3 and Finch

Both singletons display a pairwise GCS value of 22.33% with one another but share GCS values of less than 3% with all other *Rhodococcus* phages (Figure 3). These phages were isolated on different host species, with E3 isolated on *Rhodococcus equi* NCIMB 10027 and Finch isolated on *Rhodococcus erythropolis* RIA 643 (Supplemental Table 1). Both phages are of the *Myoviridae* morphotype, which has not been observed for the other sequenced and imaged *Rhodococcus* phages. The genome lengths of E3 and Finch are similar to one another (142,536 bp and 138,896 bp, respectively) with their differences attributable to discrepancies in their number of ORFs (209 and 228, respectively) (Supplemental Table 1). There is some discrepancy between the G+C% content of E3 and Finch (67.5% and 63.1%); however, these values are similar to that of their respective host’s genomes (Delegan et al., 2019; Khairy et al., 2016; Song et al., 2022; Strnad et al., 2014; Yoshida et al., 2019).

While these phages share a small portion of their genes, they show less than 75% sequence similarity to one another across their entire genomes, with most stretches showing a complete lack of nucleotide sequence similarity (not shown). For both phages, their genomes are primarily rightwards transcribed with a few small blocks of genes causing a temporary switch in their transcription orientation. E3’s genome architecture is unique in that there is a region spanning roughly 3,000 bp between genes E3_69 and E3_70 where no genes are present. This stretch of E3’s genome is not characterized by a switch in transcription orientation. Genes shared between E3 and Finch belong to phams composed primarily of *Mycobacterium* and *Gordonia* phage genes. Roughly 77% of the phams shared between E3 and Finch are present in Cluster C phages, a group of phages that infect *Mycobacterium.* Neither of their genomes code for the functions of integrase and repressor genes which implies that these phages are likely lytic. For both E3 and Finch, approximately 58% of their genes belong to orphams.

#### Singletons ChewyVIII, DocB7, Jace, Pine5, REQ1, REQ3, RRH1 and Sleepyhead

Phages ChewyVIII, DocB7, Jace, Pine5, REQ1, REQ3, RRH1 and Sleepyhead are all singletons that have pairwise GCS values between 0-18.9% with all other sequenced *Rhodococcus* phages. Most of these phages are of the *Siphoviridae* morphotype; however, the morphotypes of ChewyVIII, Jace and Sleepyhead are undetermined due to a lack of available transmission electron microscopy (TEM) images (Supplemental Table 1).

ChewyVIII was isolated on *Rhodococcus erythropolis* RIA 643 and has a genome length of 69,165 bp (Supplemental Table 1). Nearly half of its genes belong to orphams. Across its entire genome length, there are multiple points where it alternates between blocks of rightwards and leftwards transcribed genes (not shown). ChewyVIII codes for a gene with the proposed function of OCR-like antirestriction protein (ChewyVIII_86) that belongs to a pham exclusively composed of phages that infect *Streptomyces.*

DocB7 was isolated on *Rhodococcus equi* HDP1C and has a genome length of 75,772 bp (Supplemental Table 1). Roughly 60% of its genes are orphams with a majority of them localized to the right half of its genome (not shown). The first half of its genome is rightwards transcribed while the second half of its genome is leftwards transcribed. DocB7 codes for a gene with the potential function of integrase (DocB7_59); however, a repressor gene has not been identified/annotated in its genome. Given the large portion of orphams present in DocB7’s genome, more information is needed to determine whether or not this phage is likely to be temperate.

Jace was isolated on *Rhodococcus erythropolis* RIA 643 and has a genome length of 53,912 bp (Supplemental Table 1). Jace’s genome has a G+C% content of 67%, which aligns closely with that of its host type (Delegan et al., 2019; Khairy et al., 2016; Strnad et al., 2014; Yoshida et al., 2019). Approximately 50% of Jace’s genes belong to orphams. Most of Jace’s genome is rightwards transcribed; however, there are several brief switches in transcription orientation that occur throughout its genome (not shown). These switches in orientation are usually followed by only about 1-3 genes before switching back to the previous transcription orientation. The left arm of Jace’s genome primarily codes for virion structural genes (e.g., major tail subunit, major capsid protein, and minor tail protein) while the right arm mostly codes for assembly or lysis genes (e.g., DNA polymerase subunits, phosphodiesterase, and lysin B). Jace codes for the functions of both an integrase and repressor, which suggests that it is likely a temperate phage.

Pine5 was isolated on *Rhodococcus equi* 05-305 and has a genome length of 59,231 bp (Supplemental Table 1). Similar to DocB7, about 60% of its genes belong to orphams with most of them localized to the right arm of its genome (not shown). Throughout Pine5’s genome, there are multiple points where transcription alternates between blocks of rightwards and leftwards transcribed genes; in some cases these blocks can be as short as 1 or 2 genes. The majority of the genes it codes have no hypothesized functions associated with them. Of the limited number of genes with proposed functions, Pine5 does contain a gene associated with the function of lysin A (Pine5_40). The gene, Pine5_40, belongs to a pham where 86% of the other genes belong to Cluster CA phages and 11% to *Corynebacterium* phages.

REQ1 was isolated on *Rhodococcus equi* Requ28 and has a genome length of 51,342 bp (Supplemental Table 1). Approximately 84% of REQ1’s gene content is accounted for by orphams: a considerable proportion relative to the other *Rhodococcus* phages. In general, the first quarter of its genome is leftwards transcribed while the later portion is rightwards transcribed (not shown). However, there are several points across its genome where the direction of transcription switches temporarily for a small block, usually containing only 1 or 2 genes. Despite having very few genes associated with an hypothesized functional call, REQ1 does have a gene associated with the function of lysin (REQ1_83) that belongs to a pham made of genes from 2 other *Rhodococcus* singletons and 7 *Gordonia* phages.

REQ3 was isolated on *Rhodococcus equi* Requ28 and has a genome length of 39,474 bp (Supplemental Table 1). Singleton REQ3 has a G+C% content value of 65.9%, which is relatively similar to the characteristic G+C% content of its host type’s genome (Song et al., 2022). Approximately 50% of REQ3’s gene content is accounted for by orphams. The leftmost region of its genome is leftwards transcribed for 5 genes before switching to a rightwards transcription orientation for the rest of its genome (not shown). However, a single brief switch in orientation does occur in the right arm of its genome, but this temporary switch in orientation only spans a single gene. Most of the genes coding for virion structural proteins (e.g., minor capsid protein, major capsid protein, and putative tail fiber) are localized to the middle of REQ3’s genome while those genes coding for assembly/lysis proteins (e.g., lysin, endonuclease, and integrase) are located at either ends of its genome. REQ3 does code for a gene with the potential function of integrase (REQ3_01); however, a repressor gene has not been identified/annotated in its genome. Given the large portion of orphams present in REQ3’s genome, more information is needed to determine whether or not this phage is likely to be temperate.

RRH1 is the only phage in this analysis to have been isolated on the host type of *Rhodococcus rhodochrous* Rrho39 (Supplemental Table 1). This singleton has the smallest genome length out of all the sequenced *Rhodococcus* phages – just 14,270 bp – which is even small relative to the general actinobacteriophage population (Tse et al., 2022). In addition to having the smallest genome length, RRH1 has a G+C% content of 68.4%, which is the highest G+C% content value out of all the other sequenced *Rhodococcus* phages (Supplemental Table 1, Figure 5). Its genome contains an ORF of only 20 genes and is entirely rightwards transcribed (not shown). 7 of these 20 genes belong to orphams. Singleton RRH1 has 11 genes that belong to phams composed entirely of phages that infect host species other than *Rhodococcus,such* as *Gordonia, Microbacterium, Arthrobacter* or *Mycobacterium.* The only genes present in RRH1’s genome that belong to phams with gene members from at least one other *Rhodococcus* phage are RRH1_2 (terminase large subunit) and RRH1_10 (putative tail protein).

Sleepyhead was isolated on *Rhodococcus erythropolis* NRRL B-1574 and has a genome length of 43,943 bp (Supplemental Table 1). A little under 50% of Sleepyhead’s genes belong to orphams. The ends of Sleepyhead’s genome are rightwards transcribed with a block of genes that are leftwards transcribed located in the middle of its genome separating either end (not shown). In this block of leftwards transcribed genes, Sleepyhead contains genes coding for the functions of integrase and repressor (Sleepyhead_38 and Sleepyhead_39, respectively), suggesting that Sleepyhead is likely a temperate phage.

### *Rhodococcus* phage genometrics

*Rhodococcus* phage genome sizes vary from 14,270 bp (RRH1) to 142,586 bp (NiceHouse) with their G+C% content ranging from 41.2% (Grayson) to 68.4% (RRH1) (Supplemental Table 1, Figure 5). *Rhodococcus* phages have a variety of different genome end types; most end in 3’ sticky overhangs ranging from 9 to 11 bp while a smaller subset end in either direct terminal repeats ranging from 2,900 bp (Grayson) to 5,291 bp (NiceHouse and Trina) or circularly permuted genome ends. The type of genome ends for phages REQ1, REQ2 and REQ3 were not reported by the phage discoverer.

The open reading frames (ORFs) of these *Rhodococcus* phage genomes range from 20 (RRH1) to 293 (Weasles2) (Supplemental Table 1). While nearly 50% of these phages have been reported as producing clear plaques, 41 of the 58 sequenced *Rhodococcus* phages contain both an integrase gene and a repressor gene in their genome, suggesting these phages may be temperate (Hatfull, 2020). The rest of the sequenced *Rhodococcus* phages appear to lack one or more of this pair of genes, which indicates that these phages are likely lytic.

Across the 4 *Rhodococcus* clusters, there is great diversity in both the average genome size (bp) and average G+C% content of each cluster (Figure 5). A closer examination reveals a general trend between these two distinct features. Cluster CE phages display the largest average genome size relative to other *Rhodococcus* clusters with a length of 140,924 bp. Cluster CB and Cluster CC phages follow this with average genome sizes of 113,260 bp and 77,431 bp, respectively. Cluster CA phages showcase the smallest average genome size out of all the *Rhodococcus* clusters with a length of 46,597 bp. However, a reciprocal relationship appears to exist between the *Rhodococcus* clusters when it comes to their average G+C% content. Cluster CA phages display the highest average G+C% content of all the *Rhodococcus* clusters with a value of 58.8%. Following this, Cluster CC and Cluster CE phages exhibit average G+C% contents of 53.4% and 44.8%, respectively. Cluster CB phages show the smallest average G+C% content out of all the *Rhodococcus* clusters with a value of 41.3%. Therefore, there appears to be a reciprocal relationship between average genome size and average G+C% content when it comes to the *Rhodococcus* clusters (Figure 5).

For 3 of the 4 *Rhodococcus* clusters, their average G+C% content values are considerably lower than those reported for their respective host types. Clusters CB (41.3%) and CE (44.8%) both contain phages that infect *R. erythropolis* whose average G+C% content has been reported as usually exceeding 60% (Delegan et al., 2019; Khairy et al., 2016; Strnad et al., 2014; Yoshida et al., 2019). A similar finding exists between Cluster CC phages (53.4%) and their respective host type of *R. equi* whose average G+C% content values hover around 68.72% (Song et al., 2022). This discrepancy between the G+C% values of the clusters and their respective host types suggests that these phages may have only recently acquired the ability to infect these hosts in their evolutionary timeline (Pope et al., 2014). Compared to the other *Rhodococcus* clusters capable of infecting *R. erythropolis,* the average G+C% content of Cluster CA (58.8%) is closer to that of their shared host type of *R. erythropolis.*

### Evolutionary relationships relative to other actinobacteriophages

In addition to characterizing the genomic diversity of the *Rhodococcus* phages relative to one another, we asked whether the *Rhodococcus* phage community showed any similarities with phages infecting other Actinobacterial hosts. In the interest of highlighting the most meaningful relationships, only those *non-Rhodococcus* phages sharing GCS values of above 15% with at least one *Rhodococcus* phage were focused on for further analysis. A SplitsTree model was created to provide a visualization of these identified relationships between the *Rhodococcus* phages and other representative phages that infect either *Gordonia, Streptomyces* or *Arthrobacter* (Figure 1).

In a previous investigation, Bonilla et al. (2017) found that *Rhodococcus* phages in Cluster CA shared over 35% of their gene products with *Mycobacterium* phages in Cluster A. Similar to the findings from Bonilla et al. (2017), CC phages shared pairwise GCS values of 38.6% or higher with all 28 *Gordonia* phages in cluster DJ, hovering just above the current clustering threshold of 35% shared gene content (Pope et al., 2017). This finding underscores the arbitrary and sometimes limited nature of the clustering process that can sometimes omit intercluster phage relatedness such as that between this selection of *Rhodococcus* and *Gordonia* phages (Hatfull et al., 2010). The phams shared between Cluster CC and DJ phages can be found along the entire length of their aligned genomes (Figure 6). A pham containing genes associated with the function of tape measure protein (TMP), Pepy_035 and Duffington_32, is shared between Cluster CC and DJ phages. In the context of *Mycobacterium* phages, the TMP gene has been suggested as a candidate gene for cluster and subcluster assignment predictions using single-gene analysis (Smith et al., 2013). Our finding suggests that the TMP gene might not be the best candidate for assigning phages to clusters in other hosts.

**Fig 6.**
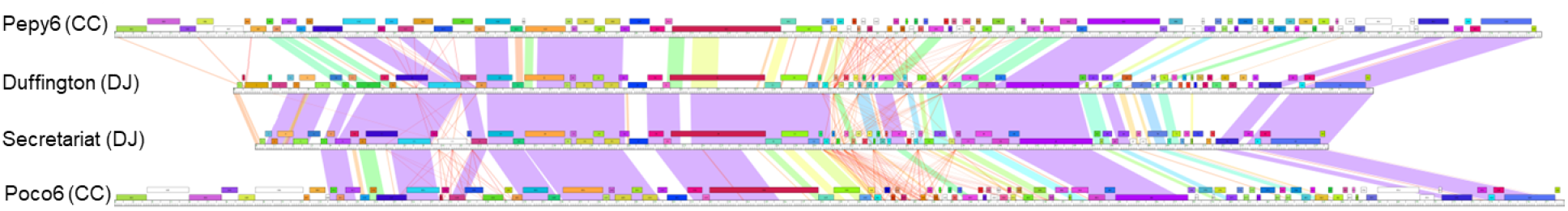
Genome comparison of representative *Rhodococcus* phages of Cluster CC and *Gordonia* phages of Cluster DJ. Genome maps were made using Phamerator. Full genome comparison of phages Pepy6 (CC), Duffington (DJ), Secretariat (DJ) and Poco6 (CC) reveals shared phams across the entire length of their aligned genomes. Cluster DJ phages Duffington and Secretariat were used for analysis since they shared the highest and lowest GCS values with both Cluster CC phages Pepy6 and Poco6. Each white ruler corresponds to an individual phage genome with the genes represented by colored boxes. Each gene’s color indicates its gene phamily (pham); white boxes represent orphams. The placement of the genes above and below the white rulers indicates rightwards and leftwards transcription, respectively. The shading between phages represents the degree of pairwise nucleotide sequence similarity between two phage genomes with purple indicating the highest similarity and red indicating lowest similarity.

Cluster CC and CE phages were found to be similar, to a lower degree, to other clusters of phages that infect other Actinobacterial hosts such as *Streptomyces* and *Arthrobacter.* Cluster CC phages Pepy6 and Poco6 were found to share GCS values exceeding 28.1% with phages from BI *(Streptomyces)* and AU *(Arthrobacter).* Cluster CE phages NiceHouse and Trina shared GCS values exceeding 16.3% with phages from Clusters BK (*Streptomyces*) and BE (*Streptomyces*); however, neither of these relationships were found to exceed the 35% clustering threshold. While Cluster CB phages (Grayson, Peregrin and Weasles2) did not share GCS values exceeding 15% with phages that infect other Actinobacterial hosts, they did share GCS values ranging from 12.1-13.2% with phages from Clusters BE and BK, both composed of *Streptomyces* phages.

### Concluding Remarks

In this investigation, we briefly summarized the isolation of 126 phages on 4 different species of *Rhodococcus: R. equi, R. globerulus, R. rhodochrous* and *R. erythropolis.* More specifically, we performed various different bioinformatic analyses on all 57 *Rhodococcus* phages that had been sequenced and annotated as of November 2022 to provide a general overview of the *Rhodococcus* phage population. Our investigation primarily focused on characterizing the intracluster and intercluster relationships of the 4 *Rhodococcus* clusters and 13 *Rhodococcus* singletons.

This research found high levels of intracluster similarity and low levels of intercluster similarity across the *Rhodococcus* phages. While these *Rhodococcus* phages display a relatively diverse genomic landscape, the discrete nature of their clusters in comparison to the populations of *Mycobacterium* phages, *Gordonia* phages and *Microbacterium* phages is likely attributable to the limited sample size of these *Rhodococcus* phages (Jacobs-Sera et al., 2020; Pope et al., 2015, 2017). Since most of these clusters, excluding Cluster CA, are composed of either 2 or 3 phages, it is likely that the findings outlined in this paper may not be entirely representative of the genomic diversity characteristic of *Rhodococcus* phages and their clusters. A continued commitment to the isolation of more *Rhodococcus* phages will most likely be required to provide a better picture of these phages and their diversity.

## Acknowledgments

We are grateful for all students in the SEA-PHAGES and PHIRE programs that contributed to the isolation and annotation of those phages included in this paper. Additionally, we are thankful for those researchers outside either program that provided similar contributions. We have composed a list of all those students and researchers that contributed in this way (Supplemental File 1).

We thank the HHMI Science Education Alliance-Phage Hunters Advancing Genomics and Evolutionary Science (SEA-PHAGES) program for programmatic support. We also thank Cristelle Hugo for early analyses. This research was supported in part by the Department of Microbiology, Immunology, and Molecular Genetics and the Dean of Life Sciences Division at UCLA.

## Author Contributions

All authors conceptualized the research. D.R.G., D.D.B., J.A.B., B.E., I.L., M.G., S.C., C.H., and B.Q., performed experiments and drafted the paper. D.R.G., A.E.G-V., and A.C.F. revised the paper. A.E.G-V. and A.C.F. supervised the research.

