## Supplemental File 1 for "Characterization of Genomic Diversity In Bacteriophages Infecting *Rhodococcus*"

Characterization of Genomic Diversity In Bacteriophages Infecting *Rhodococcus*:
SEA-PHAGES, PHIRE & Independent Contributors

### SEA-PHAGES participants:

Cabrini University:

Margaret Greenhalgh
Matthew W McGauley
Morgan Sperratore

Carnegie Mellon University:

Nathalie Chen
Noel Lau
Alex Muralles
Aru Rajeevan

College of William & Mary:

Adeola Adesuyi
Sarah Belay
Andrew Corso
Christopher Debo
Juliet Downie
Ceyda Durmaz
Jacqueline Espinoza
Marion Gilliam
Milane Gooden
Rachel Hervey
Maran Ilanchezhian
Abou Kamara
Diego Alonso Lanao
Harsha Malapati
Andrew Mattei
Sarah Modlin
Sydney Moondra
Imran Sadik

Florida Gulf Coast University:

Amanda Black
Victoria Blair
Jessica Brown
Cody Crivello
Stephanie Fine
Daniel Hansen
Abigail Huelsman
Matthew Mandio
Heather McFalls
Alec Pica
Madeline Quinn
Julia Reed-Betts
Hannah Reeves
Leah Roach
Belen Rodriguez
Gustavo Romero
Brittany Sisson
Courtney Sparrow
Matthew Mandio
Heather McFalls

Morehouse College:

Anthony Moore
Egyptian

Nyack College:

Temitope Abiodun
Torie Eggers
Jordan Jansen
Kaelan Kanai
Julianna Kranes
Abbey Lawson
Maridalia Lillis
Ruby Minaya

Nyack College (cont.):

Erin O'Brien
Christal Rolling
Valentine Sanon
Sarah Souza
Anton Teetzmann
Jacqueline Washington
Jarsibet Zapata

University of Louisiana at Monroe:

Laura Beth Aulds
Elizabeth Austin
Mitchell Broadway
Emiley Bryant
Abby Carter
C. Chamberlin
Isabel Chauvin
M. Collins
Mallory Crawford
Sachi Dhakal
Austin Dicus
Baxter Flor
Sydnie Gates
Anne Marie Hancock
Jacob Harrison
Bonnie Hemphill
Joseph Humphrey
Nathaniel Inman
Jerry Jacobs
Robbie Tyler Jester
Tyler Jester
Spencer Lambert
Brittany Little
Joseph McBride
John McKinley
Justin Netherland

University of Louisiana at Monroe (cont.):

Austin Nettles
Ryan Phillips
Megan Powell
Sabnum Pudasainy
Micheal Richers
Kristen Laurel Robinson
Dustin Rousselle
Lina Sihamath
C. Waggoner
Jordan Wagner
Dezirea Williams
Peyton Zalewski

University of Maine, Honors College:

Madeline Aubry
Mykaela Blackerby
Jacquelyn Cook
Matthew Cox
Trevor Dugal
Hanna Griffin
Victoria Mei Mayers
Ewelina Nazim
Aiden Pike
Jaelin Roberts
Ezekiel Robinson
Dylan Taplin
Marc Thibodeau

University of Wisconsin-River Falls:

Patrick Anigbogu
Bailey Braaksma
Alissa Cooper
Brent Cunningham
Rick Ellingworth
Faith Vander Galien
Cody Gostovich
Kristin Graham

University of Wisconsin-River Falls (cont.):

Krista Haglund
Collin Huebel
Bradley Koch
Madeline Kosch
Eric Liesse
Jacob Link
McKenzi Lorrig
Anna Lowery
Brittany Lubich
Kristi Mahoney
Michelle Mattison
Isaak Nicklay
Miranda Rang
Conner Satterlund
Michelle Slick
Andrea Tait

Virginia Commonwealth University:

Bret Boyd
Justin Le

Wilkes University:

Christian Laing

### PHIRE participants:

University of Pittsburgh:

Cory Hayes
Varun Iyengar
Rachael Rush
Victoria Schneider
Katie Thomas

### Independent Participants:

M. Bertoli
Samantha A. Campbell
T. N. Chan-Cortes
N. D. Cohen
Zoe A. Dyson
L. Ferguson
Sophie Foley
J. J. Gill
M. Grant
Neil F. Inglis
C. Janes
K. Lange
M. Liu
C. Moore
R. C. Orchard
Steve Petrovski
Samson P. Salifu
Robert J. Seviour
Mariela Scortti
E. J. Summer
Daniel Tillett
Ana Valero-Rello
José A. Vázquez-Boland
R. Young
