## Supplemental Table 1 for "Characterization of Genomic Diversity In Bacteriophages Infecting *Rhodococcus*"

**Supplemental Table 1: Rhodococcus Phages Included in Analysis**

| Phage Name | Cluster | Isolation Host | Length (bp) | GC% | End Type | Morphotype | ORFs | Life Cycle | Accession # |
|---|---|---|---|---|---|---|---|---|---|
| Alatin | CA | *R. erythropolis* RIA 643 | 46673 | 58.7 | 3' 10-b ext | Siphoviridae | 68 | Temperate | MF324905 |
| Alpacados | CA | *R. erythropolis* RIA 643 | 46493 | 58.9 | 3' 10-b ext | Siphoviridae | 68 | Temperate | MH271291 |
| AngryOrchard | CA | *R. erythropolis* RIA 643 | 46597 | 58.8 | 3' 10-b ext | Siphoviridae | 67 | Temperate | KY549153 |
| AppleCloud | CA | *R. erythropolis* RIA 643 | 46389 | 58.7 | 3' 11-b ext | Siphoviridae | 66 | Temperate | MF324903 |
| Belenaria | CA | *R. erythropolis* RIA 643 | 46538 | 58.6 | 3' 10-b ext | Siphoviridae | 71 | Temperate | MK524495 |
| BobbyDazzler | CA | *R. erythropolis* RIA 643 | 46641 | 58.8 | 3' 10-b ext | Siphoviridae | 68 | Temperate | KY549154 |
| Bonanza | CA | *R. erythropolis* RIA 643 | 46932 | 58.8 | 3' 10-b ext | Siphoviridae | 69 | Temperate | MF537628 |
| Bradshaw | CA | *R. erythropolis* RIA 643 | 46606 | 58.6 | 3' 10-b ext | Siphoviridae | 68 | Temperate | MH271293 |
| Bryce | CA | *R. erythropolis* RIA 643 | 46347 | 58.8 | 3' 10-b ext | Siphoviridae | 68 | Temperate | MH271294 |
| CosmicSans | CA | *R. erythropolis* RIA 643 | 46596 | 58.5 | 3' 10-b ext | Siphoviridae | 69 | Temperate | KT372002 |
| Dinger | CA | *R. erythropolis* RIA 643 | 46617 | 58.8 | 3' 10-b ext | Siphoviridae | 69 | Temperate | MN945902 |
| Erik | CA | *R. erythropolis* RIA 643 | 46429 | 58.5 | 3' 10-b ext | Siphoviridae | 69 | Temperate | MH271297 |
| Espica | CA | *R. erythropolis* RIA 643 | 46537 | 58.6 | 3' 10-b ext | Siphoviridae | 71 | Temperate | MK524487 |
| Gollum | CA | *R. erythropolis* RIA 643 | 46535 | 58.6 | 3' 10-b ext | Siphoviridae | 69 | Temperate | MH271299 |
| Harlequin | CA | *R. erythropolis* RIA 643 | 46383 | 58.8 | 3' 10-b ext | Siphoviridae | 69 | Temperate | KX611788 |
| Hiro | CA | *R. erythropolis* RIA 643 | 46854 | 58.7 | 3' 10-b ext | Siphoviridae | 68 | Temperate | MF324898 |
| Jester | CA | *R. erythropolis* RIA 643 | 46314 | 58.7 | 3' 10-b ext | Siphoviridae | 68 | Temperate | MF373842 |
| Krishelle | CA | *R. erythropolis* RIA 643 | 46985 | 58.5 | 3' 10-b ext | Siphoviridae | 70 | Temperate | MF324902 |
| Lillie | CA | *R. erythropolis* RIA 643 | 46596 | 58.6 | 3' 10-b ext | Siphoviridae | 69 | Temperate | KT990218 |
| Naiad | CA | *R. erythropolis* RIA 643 | 46619 | 58.6 | 3' 10-b ext | Siphoviridae | 68 | Temperate | MF324901 |
| Nancinator | CA | *R. erythropolis* RIA 643 | 45936 | 58.6 | 3' 10-b ext | Siphoviridae | 68 | Temperate | MH271306 |
| Natosaleda | CA | *R. erythropolis* RIA 643 | 46527 | 58.6 | 3' 10-b ext | Siphoviridae | 68 | Temperate | KX550082 |
| Partridge | CA | *R. erythropolis* RIA 643 | 46962 | 58.8 | 3' 10-b ext | Siphoviridae | 69 | Temperate | KX712237 |
| PhailMary | CA | *R. erythropolis* NRRL B-1574 | 45614 | 58.9 | 3' 10-b ext | Unknown | 65 | Temperate | MW291027 |
| Phrankenstein | CA | *R. erythropolis* RIA 643 | 46540 | 58.6 | 3' 10-b ext | Siphoviridae | 69 | Temperate | MH271309 |
| Rasputin | CA | *R. erythropolis* RIA 643 | 46568 | 58.8 | 3' 10-b ext | Siphoviridae | 69 | Temperate | MH271311 |
| RER2 | CA | *R. erythropolis* Rery29 | 46596 | 58.6 | 3' 10-b ext | Siphoviridae | 67 | Temperate | JN116827 |
| RexFury | CA | *R. erythropolis* RIA 643 | 46627 | 58.6 | 3' 10-b ext | Siphoviridae | 68 | Temperate | MF324904 |
| RGL3 | CA | *R. globerulus* Rglo35 | 48072 | 62.7 | 3' 10-b ext | Siphoviridae | 66 | Temperate | JN116826 |
| Rhodalysa | CA | *R. erythropolis* RIA 643 | 46596 | 58.5 | 3' 10-b ext | Siphoviridae | 69 | Temperate | KT375356 |
| Shuman | CA | *R. erythropolis* NRRL B-1574 | 46544 | 58.6 | 3' 10-b ext | Unknown | 70 | Temperate | MH316569 |
| StCroix | CA | *R. erythropolis* RIA 643 | 46619 | 58.6 | 3' 10-b ext | Siphoviridae | 68 | Temperate | MF324900 |
| Swann | CA | *R. erythropolis* RIA 643 | 46596 | 58.6 | 3' 10-b ext | Unknown | 69 | Temperate | MH271314 |
| Takoda | CA | *R. erythropolis* RIA 643 | 46807 | 58.7 | 3' 10-b ext | Siphoviridae | 70 | Temperate | MH271315 |
| TWAMP | CA | *R. erythropolis* RIA 643 | 46596 | 58.5 | 3' 10-b ext | Siphoviridae | 69 | Temperate | KT959213 |
| UhSalsa | CA | *R. erythropolis* RIA 643 | 46539 | 58.6 | 3' 10-b ext | Siphoviridae | 69 | Temperate | MH271319 |
| Yogi | CA | *R. erythropolis* RIA 643 | 46930 | 58.8 | 3' 10-b ext | Siphoviridae | 69 | Temperate | KX712236 |
| Yoncess | CA | *R. erythropolis* RIA 643 | 46353 | 58.8 | 3' 10-b ext | Siphoviridae | 68 | Temperate | MF189179 |
| Grayson | CB | *R. erythropolis* RIA 643 | 131801 | 41.2 | 2900bp DR | Siphoviridae | 290 | Lytic | MH153812 |
| Peregrin | CB | *R. erythropolis* RIA 643 | 133006 | 41.4 | 2937bp DR | Siphoviridae | 287 | Lytic | MH153807 |
| Weasels2 | CB | *R. erythropolis* RIA 643 | 134973 | 41.3 | 3268bp DR | Siphoviridae | 293 | Lytic | KX774321 |
| Pepy6 | CC | *R. equi* 05-306 | 76797 | 53.4 | 3' 9-b ext | Siphoviridae | 107 | Lytic | GU580941 |
| Poco6 | CC | *R. equi* MillB | 78064 | 53.3 | 3' 9-b ext | Siphoviridae | 107 | Lytic | GU580942 |
| NiceHouse | CE | *R. erythropolis* NRRL B-1574 | 142586 | 44.9 | 5291bp DR | Siphoviridae | 291 | Lytic | MT521992 |
| Trina | CE | *R. erythropolis* RIA 643 | 139262 | 44.7 | 5291bp DR | Unknown | 285 | Lytic | MF668286 |
| ChewyVIII | Singleton | *R. erythropolis* RIA 643 | 69165 | 61.8 | 3' 10-b ext | Unknown | 95 | Lytic | KX557288 |
| DocB7 | Singleton | *R. equi* HDP1C | 75772 | 56.7 | Cir Per | Siphoviridae | 105 | Temperate | GU580940 |
| E3 | Singleton | *R. equi* NCIMB 10027 | 142563 | 67.5 | Cir Per | Myoviridae | 209 | Lytic | HM114277 |
| Finch | Singleton | *R. erythropolis* RIA 643 | 138896 | 63.1 | Cir Per | Myoviridae | 228 | Lytic | MG962366 |
| Jace | Singleton | *R. erythropolis* RIA 643 | 53912 | 67 | Cir Per | Unknown | 93 | Temperate | MH153804 |
| Pine5 | Singleton | *R. equi* 05-305 | 59231 | 67.1 | Cir Per | Siphoviridae | 84 | Lytic | GU580943 |
| REQ1 | Singleton | *R. equi* Requ28 | 51342 | 66.3 | Unknown | Siphoviridae | 85 | Lytic | JN116825 |
| REQ2 | Singleton | *R. equi* Requ28 | 49330 | 65.4 | Unknown | Siphoviridae | 82 | Temperate | JN116823 |
| REQ3 | Singleton | *R. equi* Requ28 | 39474 | 65.9 | Unknown | Siphoviridae | 60 | Temperate | JN116824 |
| RRH1 | Singleton | *R. rhodochrous* Rrho39 (DSMZ43241) | 14270 | 68.4 | Cir Per | Siphoviridae | 20 | Lytic | JN116822 |
| Sleepyhead | Singleton | *R. erythropolis* NRRL B-1574 | 43943 | 61 | 3' 9-b ext | Unknown | 67 | Temperate | MK967380 |
| Whack | Singleton | *R. erythropolis* NRRL B-1574 | 49660 | 61.9 | 3' 11-b ext | Siphoviridae | 77 | Temperate | MK967393 |
